# A live yeast supplementation to gestating ewes improves bioactive molecules composition in colostrum with no impact on its bacterial composition and beneficially affects immune status of the offspring

**DOI:** 10.1101/2021.10.14.464371

**Authors:** Lysiane Dunière, Justin B Renaud, Michael A Steele, Caroline S Achard, Evelyne Forano, Frédérique Chaucheyras-Durand

## Abstract

Colostrum quality is of paramount importance in the management of optimal ruminant growth and infectious disease prevention in early life. Live yeast supplementation effect during the last month of gestation was evaluated on ewes’ colostrum composition. Two groups of ewes (n=14) carrying twin lambs were constituted and twins were separated into groups (mothered or artificially-fed) 12h after birth. Nutrient, oligosaccharides (OS), IgG and lactoferrin concentrations were measured over 72h after lambing, and bacterial community was described in colostrum collected at parturition (T0). Immune passive transfer was evaluated through IgG measurement in lamb serum. In both groups, colostral nutrient, OS concentrations and IgG concentrations in colostrum and lamb serum decreased over time, (p < 0.01) except for lactose, which slightly increased (p < 0.001) and lactoferrin which remained stable. Bacterial population was stable over time with high relative abundances of Aerococcaceae, Corynebacteriaceae, Moraxellaceae and Staphylococcaceae in T0-colostrum. No effect of supplementation was observed in nutrient and lactoferrin concentrations. In supplemented ewes, colostral IgG level was higher at T0 and a higher level of serum IgG was observed in lambs born from supplemented mothers and artificially-fed, while no effect of supplementation was observed in the mothered lambs groups. Using a metabolomic approach, we showed that supplementation affected OS composition with significantly higher levels of colostral Neu-5Gc compounds up to 5h after birth. No effect of supplementation was observed on bacterial composition. Our data suggest that live yeast supplementation offsets the negative impact of early separation and incomplete colostrum feeding in neonate lambs.

**Graphical abstract:** 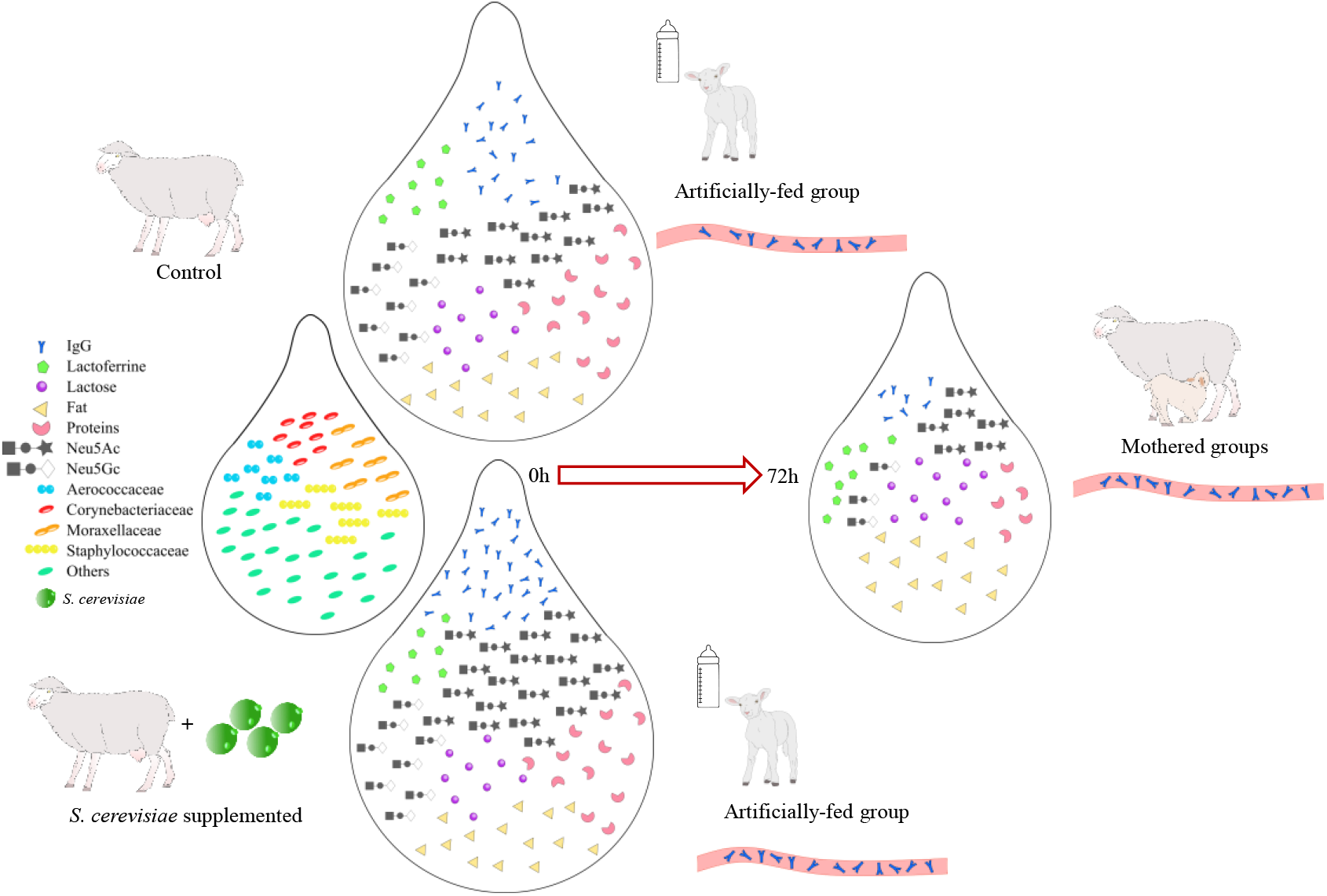

## Introduction

In ruminant industries (dairy and meat production), major economic losses occur due to diseases in the first 3 weeks of age ^1^. At birth, ruminants are hypoglycemic and agammaglobulinemic, and adequate colostrum intake is crucial for survival and control of infectious diseases in the newborn ^2^. Colostrum is the first source of nutrition in neonate ruminants, providing essential components, such as energy and nutrients, but also hormones and growth factors. The high levels of fat and proteins observed in colostrum lead to better glucose regulation in colostrum-fed ruminants (vs formula-fed animals) and improve their growth in early life ^3^. Furthermore, colostrum ingestion is of paramount importance for the neonate as it promotes Immunoglobulin (Ig) transfer from the dam to the newborn (known as Immune Passive Transfer, IPT) and provides protection against infections ^4^. Inadequate absorption of colostral Igs through the gut is known as failure of passive transfer (FPT) and is characterized by a serum IgG level lower than 10 g/L ^5^. FTP in neonatal lambs has been shown to increase neonatal mortality, with associated economical losses ^6–8^. Besides Igs, colostrum contains many other molecules involved in immunity such as lactoferrin, mucin, α-lactalbumin or serum amyloid A ^4^. Colostrum also contains leucocytes in variable concentrations ^9,10^ and high levels of oligosaccharides (OS) compared to milk. OS are considered prebiotics, i.e. highly branched sugars not digested directly by the host but by its gut microbiota. Due to its composition, colostrum ingestion immediately after birth may shape the initial digestive microbial colonization and supports gastrointestinal tract (GIT) maturation. Indeed, over the 12 first hours of life, a higher abundance of total bacteria was observed in the small intestine and colon of neonate calves, with an increase in *Bifidobacterium* genus and butyrate producers in the colon (*Clostridium* cluster XIVa) and a decrease in *E. coli* abundance ^11,12^. OS have been shown to promote *Bifidobacterium* growth ^13^. Sialylated OS are able to adhere to the intestinal epithelial cells ^14^ and increase mucosa-associated *Bifidobacterium* in the distal jejunum and colon of colostrum fed calves ^12,15^. Moreover, colostral OS have a role in host-defense by pathogen binding ^16^. Colostrum ingestion also increases GIT development by increasing cell renewal and villus development and enhancing metabolic changes to accelerate GIT maturation ^17^. Elevated levels of growth hormone, insulin, insulin growth factor and cholesterol in colostrum are involved in somatotropic axis development in the young ruminant ^18^. The sources of microbial inoculation of the ruminant neonate GIT are still under study ^19,20^ but as colostrum is the first source of energy ingested, its microbiota will be one of the first contacts of foreign elements for the neonate’s digestive tract since its birth. Few studies have depicted the microbial composition of ruminant colostrum, and those that have mainly focused on bovine colostrum, leaving information lacking in ovines.

In sheep production, artificial rearing is a common practice for prolific breeds in which supernumerary lambs are separated from their mother at very early age (0-2 days after birth) and fed milk replacer. This practice presents several detrimental short and long term effects for the young animals ^21^ and strategies are currently being studied in order to reduce the impact of mother separation on the lamb’s immunity, growth and performance and microbial digestive establishment ^22,23^. Live yeast based feed additives have been shown to enhance stabilization of rumen microbiota during challenging periods ^24,25^. Peripartum modifications of rumen microbiota composition with an increase in milk production have been observed in dairy cows supplemented with the same live yeast used in this study ^26^. Moreover, the supplemented animals presented differential expression of genes involved in pro- and anti-inflammatory processes along the digestive tract around calving, suggesting an improvement in rumen epithelial barrier associated with supplementation ^27^. An increased inflammatory status would be detrimental for the animal, as part of the metabolizable energy will be directed towards immune reactions instead of fetus development, maintenance of body condition score and the physiological changes required for lactation onset. Colostrum quality is impacted by health condition of the gestating animal ^28–30^. Probiotic supplementation peripartum could thus be a nutritional strategy to reinforce rumen functions through stabilization of microbial populations, leading to an enhancement of the dam’s health before parturition ^26,27,31^.

In this study, we hypothesized that live yeast *S. cerevisiae* supplementation of ewes during last month of gestation will improve colostrum nutritional and immune qualities impacting the immune passive transfer to their lambs afterwards and also modify colostrum microbiota. More precisely, we investigated differences in colostrum nutrients, lactoferrin and IgG concentrations. Oligosaccharide compositions were depicted through highly specific LC-MS techniques, and bacterial population was described through qPCR and 16S rDNA sequencing approaches.

## Material and Methods

### 1 Diets and animals

The animal trial was conducted at the animal facilities of INRAE Herbipôle Experimental Unit UE1414 (Clermont Auvergne Rhône Alpes). Procedures on animals were carried out in accordance with the guidelines for animal research of the French Ministry of Agriculture and all other applicable national and European guidelines and regulations for experimentation with animals (see http://www2.vet-lyon.fr/ens/expa/acc_regl.html for details). The protocol was approved by the Regional Ethics Committee for Animal Experimentation C2EA-02 and authorized by the French Research Ministry with the reference number 14981-2018050417167566V3.

Twenty-eight gestating ewes (*Ovis aries*, Romane breed) were used for this study. All of them carried two fetuses. They were selected following echographic evaluation a few weeks before the start of the trial and assigned to two groups (**Control = C, Supplemented = SC**) which were balanced homogeneously according to age, parity, body condition score and live weight.

One month and a half before estimated date of parturition, ewes were transferred from the farm unit to a room equipped with Biocontrol CFRI systems (www.BioControl.no) which allowed to control individual concentrate intake. Ewes were identified by means of ear RFID (radio frequency identification) transponders for specific access to the manger. As the yeast supplement was incorporated into the experimental concentrate, it was important to ensure that each animal had the same quantity of concentrate ingested, and at the same time of the day. Ewes were progressively adapted to the concentrate during the month before the start of the trial, by increasing the amount of concentrate fed daily up to 800 g /d/animal. Then, until parturition, each ewe received this fixed amount of concentrate (Moulin de Massagettes, Massagettes, France, Supplementary Table S1) daily, distributed once at 8:00 am, followed by 2 kg of meadow hay. Good quality water was offered *ad libitum*.

After 2 weeks of adaptation to the BioControl system and to this diet, the two groups received their allocated experimental concentrate, the only difference being the incorporation of the live yeast product – *Saccharomyces cerevisiae* CNCM I-1077 (Levucell SC TITAN, Lallemand SAS, Blagnac, France) – for the SC group. The rate of inclusion in the concentrate was calculated to bring 8 × 10^9^ CFU per day per individual. A few days before their estimated date of parturition, ewes were transferred to a maternity unit which was split into two large pens separated by a concrete wall, to ensure that no contact could occur between the groups. Bedding was made of straw. Animals were then kept in these pens until the end of experiment.

After parturition, the concentrate was not supplemented anymore with live yeast product. Thus, live yeast supplementation to the SC group occurred only during the end of the gestation phase and was stopped at parturition. The composition of the postpartum diet was modified to meet the requirements of the dam in order to feed only one lamb (the other one was directed to an artificial milk feeding system). So, each ewe was fed with 600 g of concentrate and 3 kg of meadow hay, covering slightly more than 135 % of its energy needs.

At birth, twin lambs were kept with their mother in an individual birth pen for about 12h to ensure a first colostrum uptake. During this period the lambs were weighed and ear tagged. Animal care was performed if necessary, and lambs were trained to reach the teats. Then one of the twins was separated from the dam, entered into the Artificially fed-group (**Art-**) and was moved in another pen, whereas the other twin was kept with the dam (Mothered group, **Mot-**). The lambs were chosen on the basis of their birth weight and sex to constitute homogeneous groups. Thus, four groups of lambs were constituted, named Cont-Art, Suppl-Art, Cont-Mot, Suppl-Mot Lambs from the four groups considered in this study for blood sampling. Artificially fed lambs were fed with milk replacer (Agnodor, Bonilait Protéines, France). They had been bottle fed for the first few hours/days and as soon as they were comfortable and autonomous, they were fed with a milk bucket equipped with teats, then an automatic milk replacer feeder was put in place to offer milk *ad libitum*. The same concentrate given to the ewes was offered from the second week, along with good quality hay and good quality water, but was very weakly consumed during the first days. However, as the bedding was made of straw, lambs could also have access to fibre through this bedding area. Weaning occurred when the lambs reached about 14 kg, as it is usually done at the experimental farm.

### 2 Sample collection

#### a Colostrum sampling from ewes

Samples were taken manually and collected at lambing (T0), 5h, 12h, 24h and 72h after lambing without oxytocin injection. The samples were collected from the two teats with gloves and with prior cleaning of the teats with a clean tissue immersed in hot water. A camera was installed in each experimental room to monitor potential night lambing and further calculate the time when colostrum would be sampled. Samples were aliquoted in several Eppendorf sterile microtubes and rapidly frozen at -20°C for further analyses. Due to physiological variations in lambing time and colostrum yield, the quantity collected varied greatly among animals and time and some ewes couldn’t be sampled at each time points. The final number of colostrum samples was 12 samples at T0h (C = 6, SC = 6), 21 samples at T5h (C = 10, SC = 11), 24 samples at T12h (C = 11, SC = 13), 24 samples at T24h (C = 13, SC = 11) and 26 samples at time T72h (C = 13, SC = 13).

#### b Blood sampling from lambs

Blood samples were collected from all lambs at day 2, 7, 28 and 55 of age through the jugular vein in dry collection tubes (Becton Dickinson, Franklin Lakes, NJ, USA) by a qualified technician. Dry tubes were set for clotting for at least 1 h at room temperature, centrifuged at 4 500 rpm for 20 min at 4 °C and serum supernatants were stored at -80 °C until further analysis.

### 3 Sample analyses

#### a Biochemical analyses

Nutrient composition of colostrum was determined from samples diluted 1/10, 1/100 or 1/1000 in PBS 1X as follows: lactose content was measured from 200 μl of diluted samples with Lactose/D-Galactose (Rapid) Assay Kit (Megazyme, Wicklow, Ireland) according to the manufacturer instructions; protein content was measured from 100 μl of diluted samples with Pierce™ BCA Protein Assay Kit (ThermoFisher, Rockford, IL, USA) and lipid content was analyzed by hydrolysis according to an internal method adapted from NF ISO 6492 by Artemis Laboratory (Janzé, France) from 1 ml of frozen raw colostrum.

Bioactive molecules were analyzed as follows: lactoferrin concentration was measured from 50 μl of diluted samples through Sheep Lactoferrin (LF) ELISA Kit (Mybiosource, San Diego, CA, USA) according to the manufacturer recommendation; the Immunoglobulin G (IgG) concentration was analyzed by radial immunodiffusion by CIAL Sud Ouest laboratory (Auch, France) from 1 ml of raw colostrum and 1 ml of lamb blood sample. For biochemical analyses, no technical replicate was performed due to volume limitation, but all biological replicates were analyzed (i.e. from 6 to 14 animals per group).

#### b High-resolution LC-MS colostrum analysis

All LC-MS data was acquired with a Thermo Q-Exactive Orbitrap^®^ mass spectrometer coupled to an Agilent 1290 HPLC system. Analytes were resolved by hydrophilic liquid interaction chromatography (HILIC) with a 350 ul min^-1^ flow rate. Two microliters of sample was injected onto a PEEK lined Agilent HILIC-Z (2.1 × 100 mm, 2.7 μm; Agilent) column maintained at 35 °C. Compounds were resolved with mobile phases of 10 mM ammonium formate + 0.1% formic acid in water (A) and 10 mM ammonium formate + 0.1% formic acid in 90% acetonitrile (B) operating with the following gradient: 0 min, 90% B; 1.0 min, 90% B; 5.0 min, 62% B; 5.5 min, 30% B; 10.5 min, 30% B, 11.0 min 90% B and 15.5 min, 90% B. Sample of 100 μl of each raw colostrum sample was diluted with 400 μl of ddH_2_O. These samples were extracted using the method developed by Fischer et al. ^15^. A 5 μl spike of the internal standard (IS: β1–3-gal-N-acetyl-galactosaminyl-β1–4-gal β1–4-Glc; GalNAc, 1000 μg/ml) was added to the diluted colostrum sample, for a final IS concentration of 9.9 μg/ml. Finally, a 100 μl aliquot was diluted with 100 μl acetonitrile (0.2% FA, 20 mM ammonium formate) and placed in a 250 μl polypropylene HPLC vial prior to LC-MS analysis. The following conditions were used for heated electrospray ionization (HESI): capillary voltage 5 kV; capillary temperature, 330 °C; sheath gas, 32 arbitrary units; auxiliary gas, 10 units; probe heater temperature, 280 °C and S-Lens RF level, 50%.

##### Non-targeted chemical analysis

Non-targeted analysis was performed on a subset of samples: 12 samples at T0h (C = 6, SC = 6), 21 samples at T5h (C = 10, SC = 11) and 9 samples at T72h (C = 3, SC = 6). Samples were analyzed in both positive and negative ionization modes, at 140,000 resolution, automatic gain control (AGC) of 3×10^6^, maximum injection time (IT) of 512 ms and mass range 150 to 1250 m/z. Ten microliters of all samples were combined as a pooled QC composite sample that was analyzed at the beginning and end of the LC-MS analysis to verify minimal instrumental drift. Composite samples were also analyzed by a top 8 data-dependent acquisition for compound identification consisting of a 35,000 resolution, 3×10^6^ AGC, 128 ms max IT followed by MS/MS at 17,500 resolution, 1×10^5^ AGC, 64 ms, 1.2 m/z isolation window and 25,35 steeped collision energy. Thermo .raw files were converted to .mzml format using Protewizard ^32^ with peak peaking filter applied. Features were detected using the XCMS package ^33^ with the centWave method (ppm tolerance 1.0 ^34^). The signal to noise threshold was set to 5, noise was set to 1×10^6^ and pre-filter was set to six scans with a minimum 5,000 intensity. Retention time correction was conducted using the obiwarp method ^35^. Grouping of features was set to those present in at least 25% of all samples (retention time deviation 10 s; m/z width, 0.015). The ‘fillPeaks’ function with default settings. Remaining zeros values were imputed with two thirds the minimum value on a per mass basis. Compounds were identified by accurate mass, comparison of retention times to authentic standards or by accurate mass and also comparison of fragmentation patterns to MS/MS databases ^36^.

##### Quantification of sialyl oligosaccharides

Eight analytes (Figure 5 and Supplementary table S2) were quantified within all samples by single stage, high resolution mass spectrometry based on their accurate mass (±3 ppm). The target analytes were monitored in negative ionization mode by 2 SIM scans between mass ranges of *m/z* 625-710 and 915-940. Both SIM scans were performed at a resolution of 17,500 resolution, automatic gain control (AGC) of 3×10^6^, and maximum injection time of 64 ms. The LC method was modified for these target analytes as follows: Mobile phase B was held at 90% for 1.0 min, before decreasing to 62% over 4 min followed by a decrease to 30% over 0.5 min. Mobile phase B was held at 30% for 4 min, before returning to 90 % B over 0.5 min and equilibrated for 2 min. Given their high structural similarity and retention times, the N-acetylneuramic (Neu5Ac) containing compounds were used as surrogate standards for the quantification of the corresponding N-glycoylneuraminic (Neu5Gc) containing analytes. The recovery efficiency (R_e_%) for GalHNAc, 3’ASL, 6’ASL, 3’ASLN, 6’ASLN were calculated by fortifying 100μL composite colostrum samples with each analyte before extraction (pre-spike) and fortifying a second set after extraction. The signal suppression/enhancement (SSE%) was calculated by the signal ratio of composite colostrum samples fortified after extraction with a fortified water sample. For targeted analysis, all collected samples from T0h to T72h were processed and analyzed in duplicate.

#### C Colostrum microbiota analysis

DNA was extracted from at least 250 mg of raw colostrum using the Quick DNA Fecal/Soil Microbe kit (Zymo Research, Irvine, CA, USA). DNA yield and quality were determined after Nanodrop 1000 and Qubit spectrophotometric quantifications. DNA extracts were stored at -20 °C until analysis. An average of 20.83 ± 73.34 ng/μl of DNA were extracted from raw colostrum samples (dsDNA, Qubit quantification), except for one sample from SC group whose quality was not sufficient for further analysis, and which was thus discarded from bacterial analyses.

Total bacteria population was quantified using qPCR method, with specific primer set and PCR conditions targeting ribosomal RNA gene according to Bayat et al., ^37^. The absolute abundance of total bacteria was expressed as the number of gene copies per microgram of colostrum. The standard curves (from 10^2^ to 10^9^ copies of bacterial 16S rDNA) were prepared by serial dilutions according to Mosoni et al., ^38^. Efficiency of the qPCR for each target varied between 97 and 102 % with and a regression coefficient above 0.95.

Microbial diversity and composition of colostrum were studied in T0 samples (ie: 6 ewes per treatment group), using 16S rDNA amplicon sequencing. The hypervariable V3-V4 regions of the 16S rRNA gene were targeted for sequencing (466bp; 341F 5’-CCTAYGGGRBGCASCAG-3’; 806R 5’-GGACTACNNGGGTATCTAAT-3’). High-throughput sequencing was performed on a Illumina MiSeq sequencer by the GeT-PlaGe core facility (INRAe Transfer, Toulouse, France). MiSeq Reagent Kit v3 was used according to the manufacturer’s instruction (Illumina Inc., San Diego, CA). Bioinformatics analyses were performed using GenoToul bioinformatics facility (INRAe, Toulouse, France). Sequences were processed using FROGS 3.2 pipeline on Galaxy platform ^39^. Briefly, sequences were clustered in Operational Taxonomic Units (OTUs) using SWARM algorithm ^40^. Chimeric sequences detected by samples using UCHIME algorithm ^41^. A total of 464 858 reads were merged and processed and 437 504 sequences were kept after chimera removal. Singleton OTUs were excluded and the remaining 2 311 OTUs were affiliated with SILVA 138.1 database using BLAST algorithm (97% sequence identity threshold).

#### d Statistical analysis

The number of animals enrolled in this study was limited by the experimental facility capacity and the criteria retained for animal inclusion in each experimental group. More precisely, ewes had to be gestating with two fetuses and each experimental group was balanced by ewe’s body score condition, live weight, age and parity. According to these criteria, 14 ewes were enrolled in each experimental group. Graphical representations and statistical analyses were performed using GraphPad Prism 9.0.2. The effect of Supplementation (C or SC) and Time factors and their interaction were evaluated using mixed model with repeated time and multiple comparisons with Sidak’s adjustment. Colostrum qPCR data were log_10_ transformed before statistical analysis. For lamb serum analysis, a linear mixed model with repetitions was applied considering 3 factors: Time, Supplementation and Rearing mode (Art- or Mot-groups) and their interactions. Multiple comparisons were made according to Tukey’s test. For colostrum microbial sequencing data, Mann Whitney test was performed on relative abundance of each considered taxa. Statistical significance was determined at a p-value < 0.05 and trends discussed at p-value < 0.10. Significant effect of Supplementation factor was indicated in Figures with # p < 0.1, * p < 0.05, ** p < 0.001 and *** p < 0.0001.

## Results

### 1 Nutrient composition

The nutrient composition of colostrum was studied over the 72h after lambing (Figure 1). Fat (1a), lactose (1b) and protein (1c) concentrations varied greatly among individuals and no statistical differences were observed between Control (C) and Supplemented (SC) groups, but a strong Time effect was observed (Table 1). Fat concentration decreased over time while a slight increase of lactose concentration was observed with time. A drastic decrease of protein concentration was measured from the first hours after lambing.

**Table 1:** p-values associated to statistical analysis of nutrient composition of colostrum samples with linear mixed model.

| Nutrient (mg/ml) | Supplementation | Time | S x T |
| --- | --- | --- | --- |
| Fat | 0.82 | **0.01** | 0.15 |
| Lactose | 0.11 | **< 0.001** | 0.53 |
| Protein | 0.91 | **< 0.001** | 0.39 |

**Figure 1:**
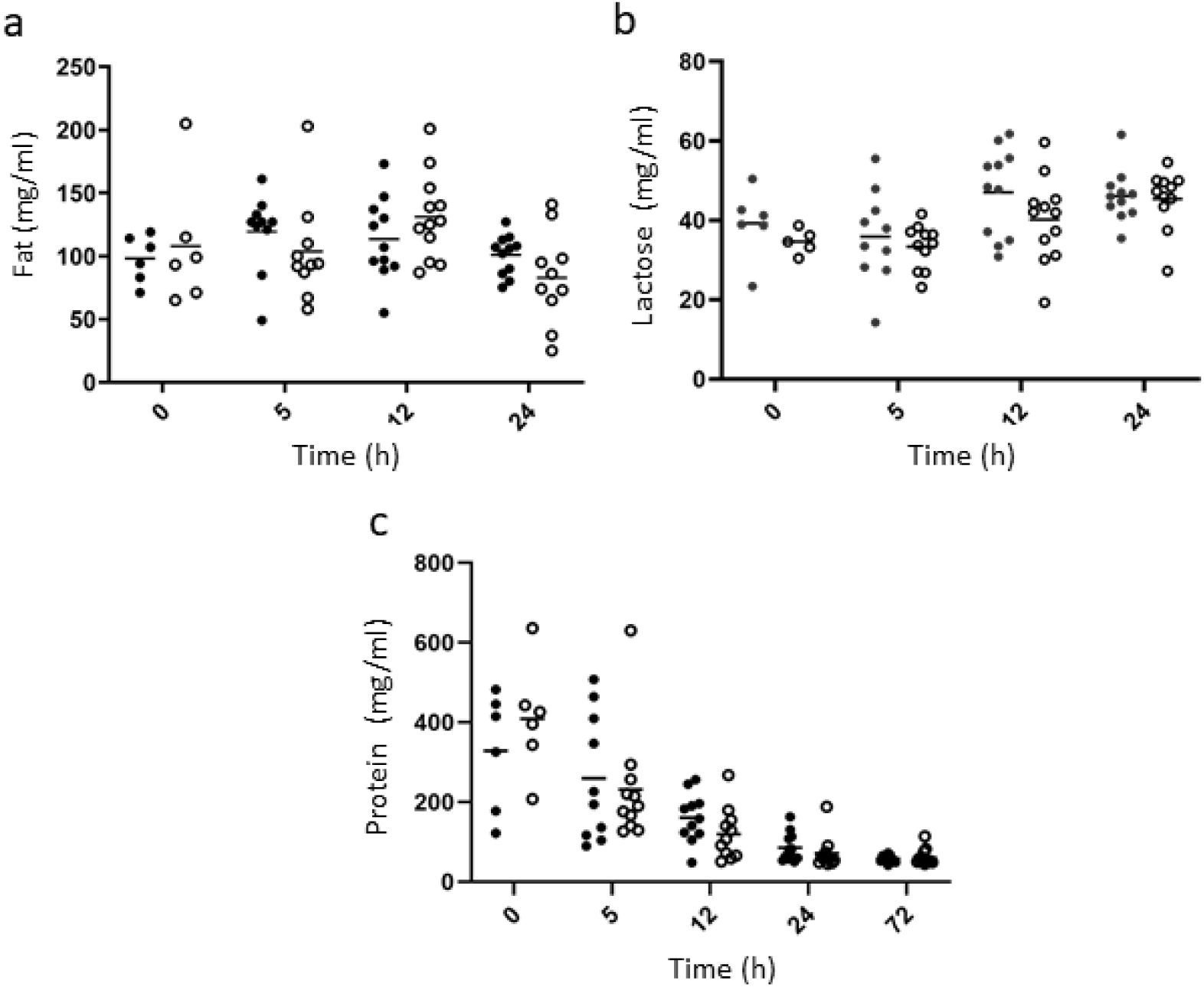
Evolution of fat (a), lactose (b) and protein (c) concentrations (mg/ml) over time in colostrum samples in C (dark circle) and SC (open circle) ewe groups.

### 2 Bioactive molecule composition

Lactoferrin concentration in colostrum was quite stable over time (Figure 2a, Table 2). Great intra-individual variations were observed among the animals, especially at early time points. Interestingly, a numerically higher level of lactoferrin (+15.8 %) was quantified in colostrum of supplemented ewes at 24h (3.79 ± 0.28 μg/ml and 4.38 ± 0.52 μg/ml in C and SC groups respectively) after lambing. Immunoglobulin G (IgG) was measured over time in colostrum (Figure 2b). A large variation in IgG concentrations was observed among animals during the first 12h post-partum. Concentrations decreased rapidly to reach low levels after 24h (from 52.89 ± 31.61 and 79.50 ± 15.24 mg/ml at 0 h to 3.73 ± 2.48 and 4.03 ± 4.24 mg/ml after 72h for C and SC groups, respectively, p < 0.001). There was a tendency for Time x Supplementation interaction and the IgG concentrations in the colostrum of supplemented ewes were higher than those of control at T0 (p = 0.046, Sidak’s multiple comparison test).

**Table 2:** p-values associated to statistical analysis of bioactive molecules’ concentrations in colostrum samples with linear mixed model.

| Bioactive molecules (mg/ml) | Supplementation | Time | S x T |
| --- | --- | --- | --- |
| Lactoferrin | 0.14 | 0.17 | 0.49 |
| IgG | 0.28 | **< 0.001** | **0.10** |

**Figure 2:**
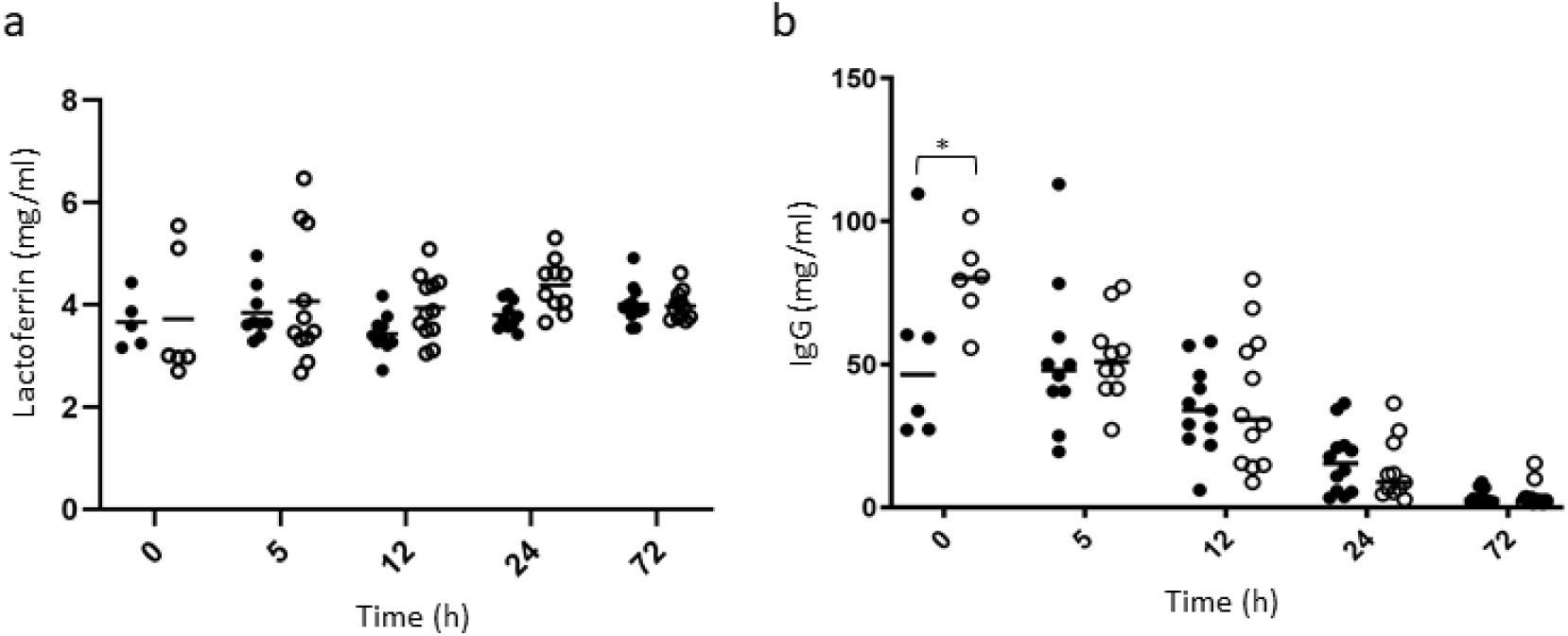
Lactoferrin (2a, mg/ml) and IgG (2b, mg/ml) concentrations over time in colostrum samples of C (dark circles) and SC (open circles) groups. Significant effects of Supplementation factor are indicated with a bracket with * p < 0.05.

IgG concentrations were also measured in the serum of lambs (Figure 3). Strong significant effects of Time, Supplementation and their interaction were observed, as well as a tendency for the interaction of the 3 factors considered (Table 3). The serum IgG concentration drastically decreased in all groups during the first 3 weeks of life from a predicted mean of 28.71 mg/ml at 2d of age to 7.42 mg/ml after 28d and remained stable up to 55d (7.01 mg/ml). Among Artificially fed lambs, a significant higher level of IgG was observed in serum of lambs born from supplemented mothers at day 2 (p < 0.001, Tukey’s multiple comparison test). Interestingly, no significant difference of IgG concentration was observed among artificially fed lambs from supplemented mothers and lambs kept with their mother whatever the supplementation status.

**Table 3:** p-values associated to statistical analysis of IgG concentrations in the serum of lambs with linear mixed model.

| IgG (mg/ml) | P-value |
| --- | --- |
| Time | **< 0.001** |
| Rearing mode | 0.28 |
| Supplementation | **0.01** |
| T x R | 0.19 |
| T x S | **0.003** |
| R x S | 0.10 |
| T x R x S | **0.09** |

**Figure 3:**
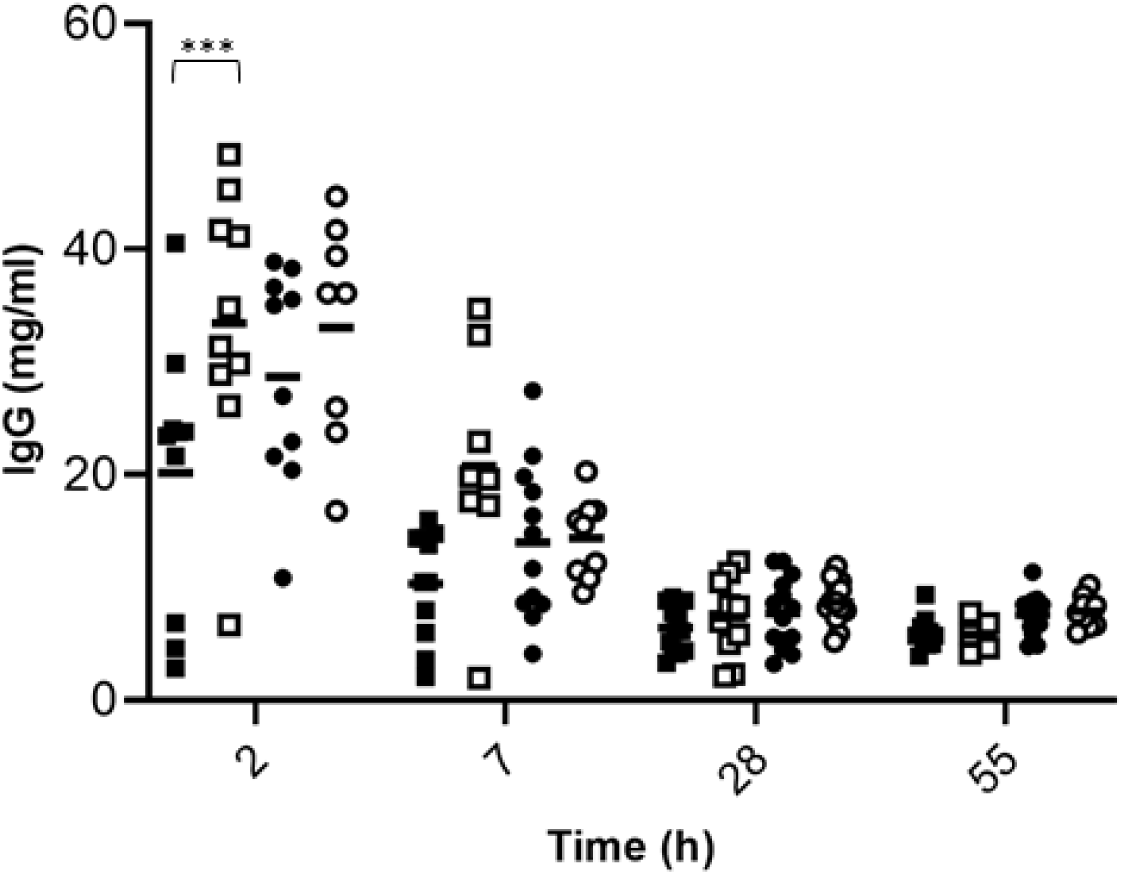
IgG concentration (mg/ml) over time in serum samples of lambs born from C (dark) or SC (white) ewes and raised with their mother (circle) or artificial fed (square). Significant effects of Supplementation factor are indicated in the graph with a bracket with *** p < 0.0001.

### 3 Metabolomic analysis of colostrum samples

#### a Non-targeted analysis

Colostrum samples from 0 h, 5 h and 72 h post-partum of both groups were analyzed in positive and negative ionization mode. In positive mode, the most intense signals corresponded to carnitine, acetyl carnitines as well as lactose and glycerophosphocholine (Figure 4a). In negative ionization mode (Figure 4b), the major analytes detected were two isobaric peaks matching the formula of lactose (α- and β-anomers of lactose). Two other major peaks eluting later in the run are uridine-diphosphate (UDP) hexose and UDP-acetylglucosamine, the precursors to lactose and *N-*acetyllactosamine respectively.

**Figure 4:**
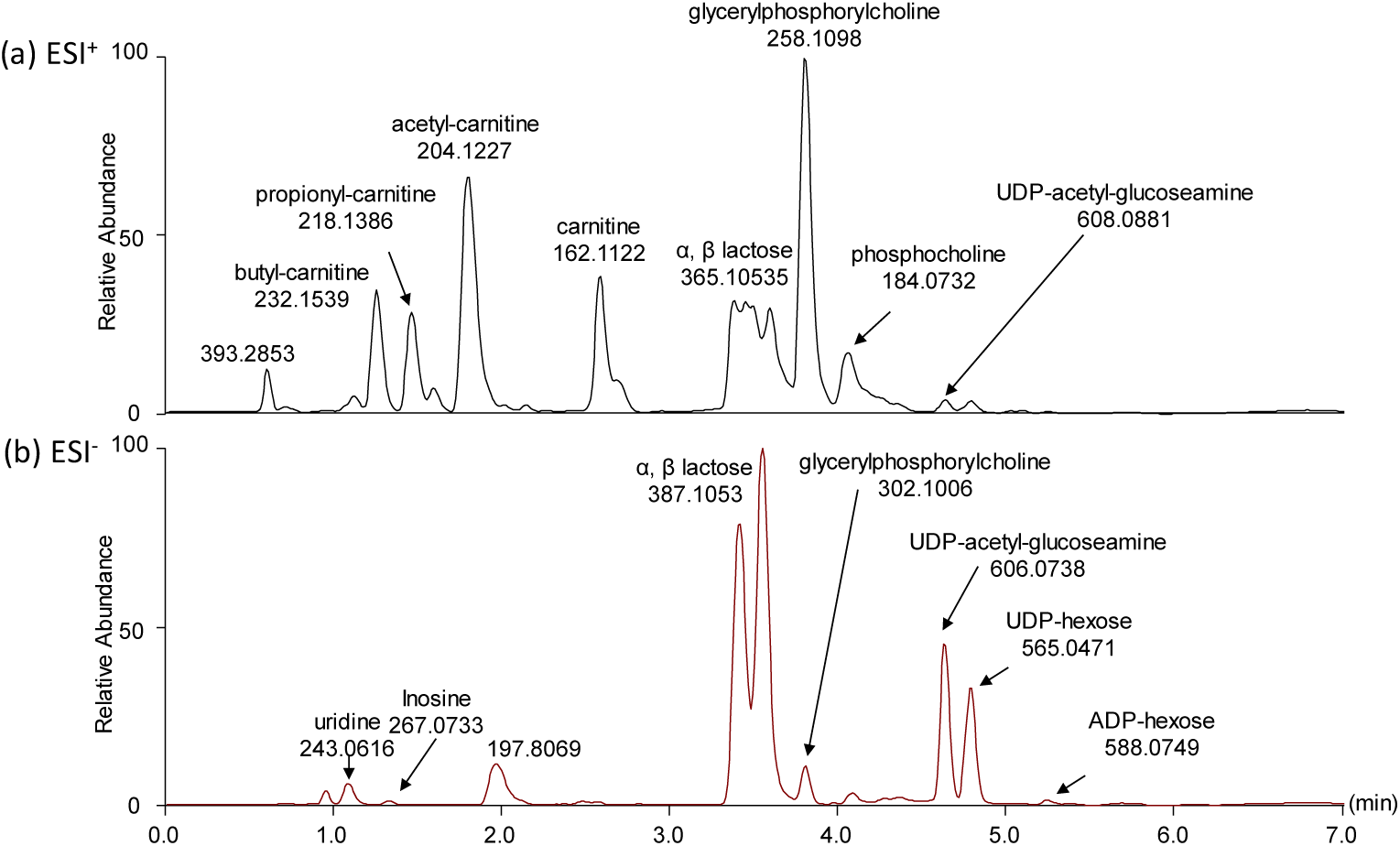
Colostrum samples analyzed in (a) positive and (b) negative ionization mode.

In positive ionization mode, 583 features were extracted among which 12, 21 and 3 features were differentially expressed (p < 0.01) at 0, 5 and 72 hours post-lambing, respectively (Supplementary table 3). Of all differentially expressed features, only 3, 4 and 1 were decreased in the SC group after 0 h, 5 h and 72 h, respectively. In negative ionization mode, fewer features were extracted at the same defined noise level. Of the 355 features extracted, only 6, 7 and 4 were differentially expressed (p < 0.01) at 0, 5, and 72 hours post-lambing. A significantly altered feature of formula C_25_H_42_N_2_O_20_ was detected in both positive and negative ionization modes. In negative ionization mode, another significantly increased feature has a putative formula of C_23_H_39_NO_19_.

The identity of these differentially expressed analytes was characterized by MS/MS using spectra of commercial standards of 6’acetyl-sialyllactose (6’-ASL) and 6’acetyl-sialyllactosamine (6’-ASLN, Supplementary Figure S1). The distinct spectra observed indicated that an additional oxygen atom is present on the sialyl residue. These compounds were thus identified as N-glycolylneuraminic (Neu5Gc) lactose and Neu5Gc lactosamine. The precursors Neu5Ac OS were also detected in high concentrations in analyzed samples (Figure 5).

**Figure 5:**
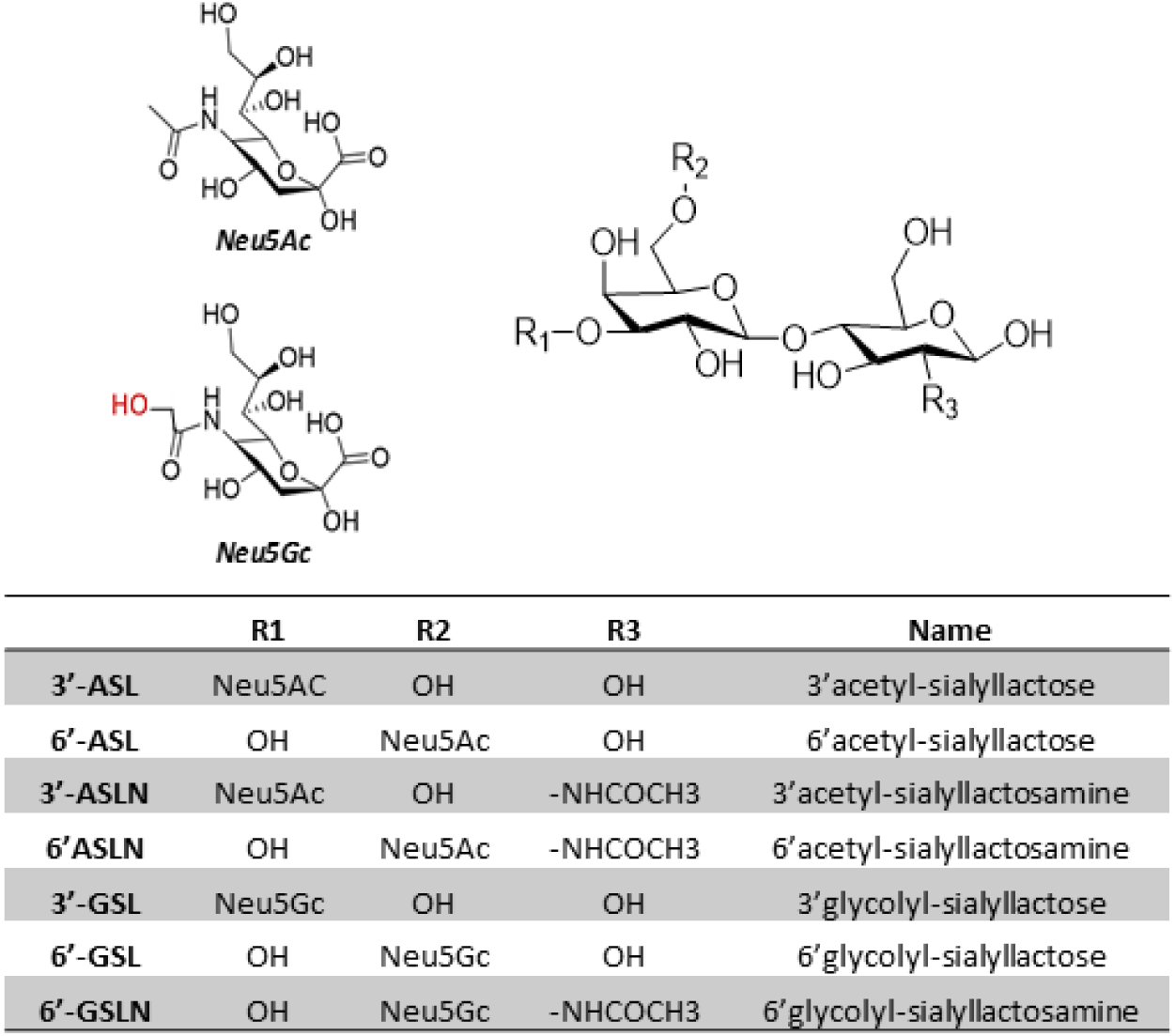
Spatial representations, name and nomenclature of the structures of major sialyl-oligosaccharides identified in ewe’s colostrum.

#### b Targeted analysis of colostrum samples

The sialyl oligosaccharides identified by non-targeted analysis were quantified by high resolution MS over 72 h post-lambing to determine if supplementation affected their production. A strong and significant Time effect was observed for total sialylated OS (Total), Neu5Ac, Neu5Gc and any of the OS structures observed (3’- and 6’-sialyllactose, disialyllactose and 3’- and 6’-sialyllactosamine, Table 4, Figure 6). No effect of Supplementation was observed among the Neu5Ac OS identified while significant Supplementation effect and interaction with Time was observed for 6’-GSL, 6’-GSLN and total Neu5Gc. More precisely, a higher concentration of Neu5Gc was observed in SC group in the T0 colostrum (p = 0.018). The concentration of 6’GSL (Figure 7a) was significantly increased as well at T0 (89.91 ± 39.57 mg/l and 147.38 ± 62.92 mg/l in C and SC groups, respectively, p = 0.042) and also 5 h after lambing (85.39 ± 43.53 mg/l and 130.07 ± 53.94 mg/l in C and SC groups, respectively, p = 0.044). Concentration of 6’GSLN in SC group was about twice that measured in C group at the same time points (119 ± 40.37 mg/l and 248.47 ± 120.09 mg/l in C and SC groups, respectively, p < 0.001 at 0 h; 106.26 ± 42.59 mg/l and 180.22 ± 70.27 mg/l in C and SC groups, respectively at 5 h; p = 0.009, Figure 7b). After 12 hours, no differences between the groups were measured.

**Table 4:** p-values associated to statistical analysis of OS concentrations in colostrum samples with linear mixed model.

| OS (mg/l) | Time | Supplementation | T x S |
| --- | --- | --- | --- |
| 3'ASL | < **0.001** | 0.89 | 0.87 |
| 6'ASL | < **0.001** | 0.14 | 0.64 |
| 3'ASLN | < **0.001** | 0.31 | 0.41 |
| 6'ASLN | < **0.001** | 0.26 | 0.63 |
| DSL | **0.007** | 0.79 | 0.75 |
| Total Neu5Ac | < **0.001** | 0.35 | 0.77 |
| 3'GSL | < **0.001** | 0.51 | 0.30 |
| 6'GSL | < **0.001** | **0.03** | **0.07** |
| 6'GSLN | < **0.001** | **0.004** | **0.001** |
| Total Neu5Gc | < **0.001** | **0.06** | **0.02** |
| Total sialyl OS | < **0.001** | 0.12 | 0.13 |

**Figure 6:**
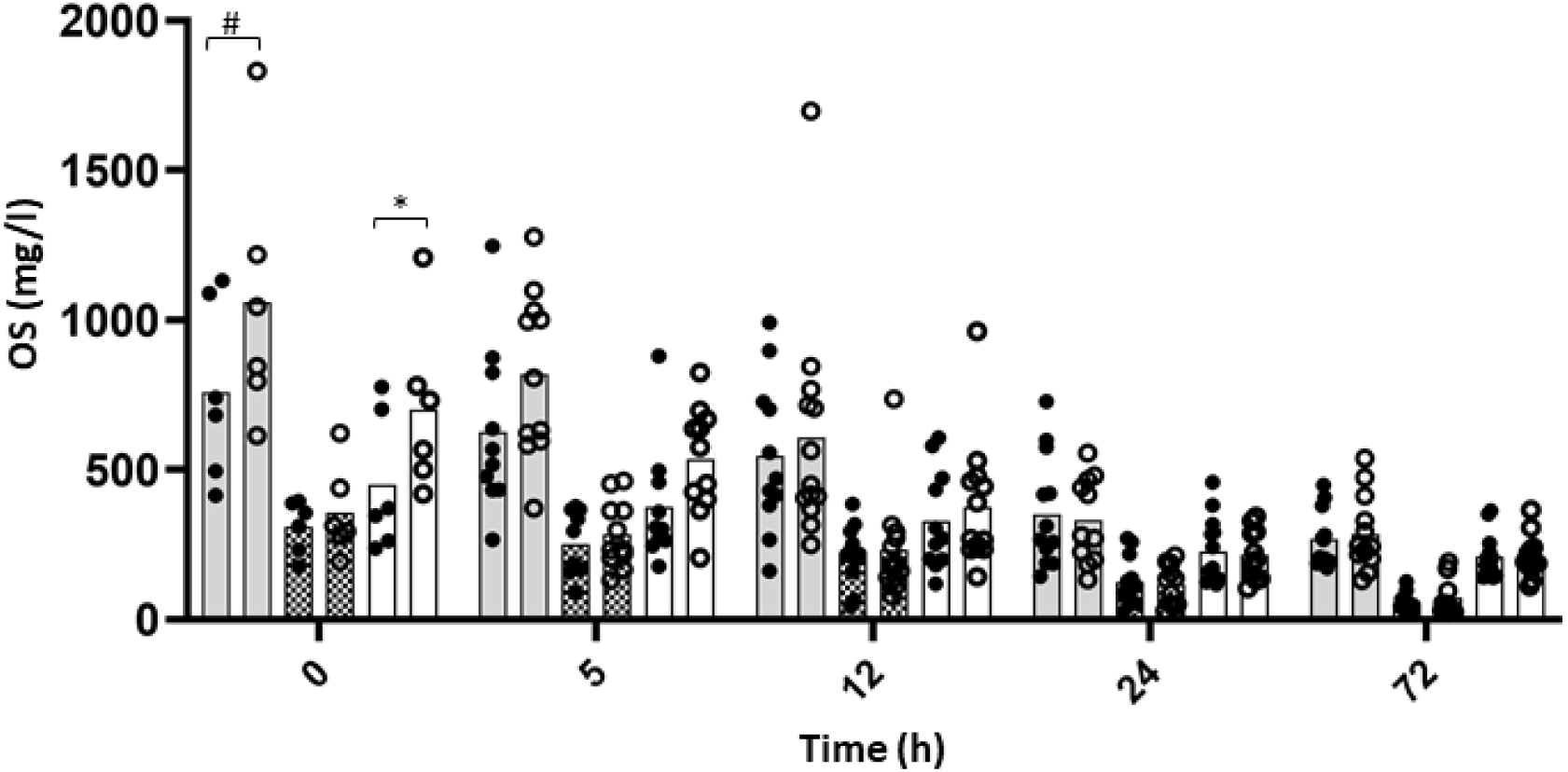
Concentrations of sialyl oligosaccharides in colostrum of C (dark circle) or SC (open circle) ewes over time. Grey, hatched and open bars represent the total sialylated OS, Neu5Ac and Neu5Gc OS content, respectively. Significant effects of Supplementation factor are indicated in the graph with a bracket with # p < 0.1, * p < 0.05, ** p < 0.01 and *** p < 0.0001.

**Figure 7:**
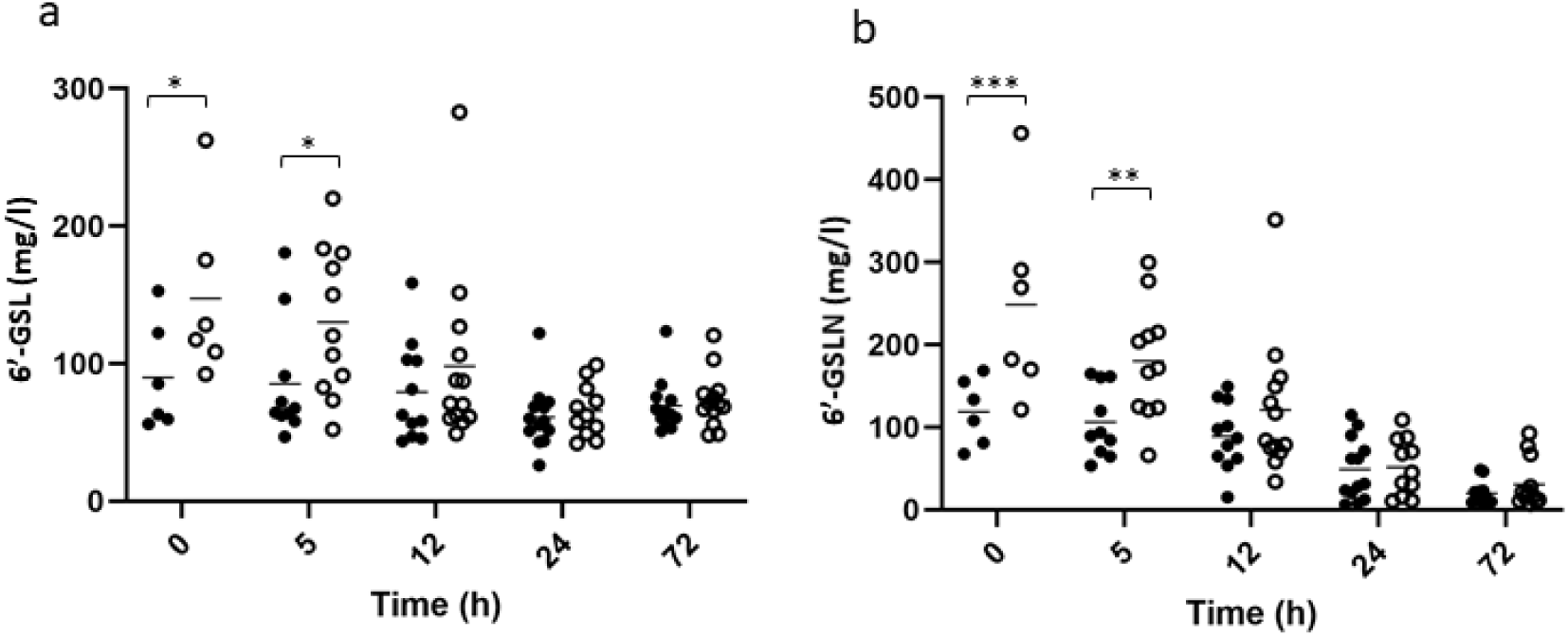
Concentrations of 6’GSL (a) and 6’-GSLN (b) OS in colostrum of C (dark circle) or SC (open circle) ewes over time. Significant effects of Supplementation factor are indicated in the graph with a bracket with # p < 0.1, * p < 0.05, ** p < 0.01 and *** p < 0.0001.

The biosynthetic pathway for both Neu5Ac OS and Neu5Gc OS has been reconstructed from the non-targeted LC-MS data (Figure 8). A Neu5Ac residue is loaded onto a cytosine-monophosphate molecule (CMP) which can transfer the Neu5Ac to the 3’ or 6’ position of lactose or *N-*acetyllactosamine. Alternatively, CMP-Neu5Ac can be hydroxylated enzymatically to CMP-Neu5Gc. The statistical analysis of the reconstructed Neu5Gc pathway was performed on colostrum samples collected 5 h pos-partum as the statistical significance was the highest at that sampling time. No significant difference was observed between C and SC groups for Neu5Ac, lactose, *N-*acetyllactosamine nor the Neu5Ac OS. CMP-Neu5Ac was also not altered by supplementation. In SC group, the elevated levels of Neu5Gc containing oligosaccharides (glycolyl-SLN and glycolyl-SL; p < 0.01) concomitant with the decreased levels in CMP-Neu5Gc (p = 0.03) suggest an increased activity of sialyltransferase enzymes.

**Figure 8:**
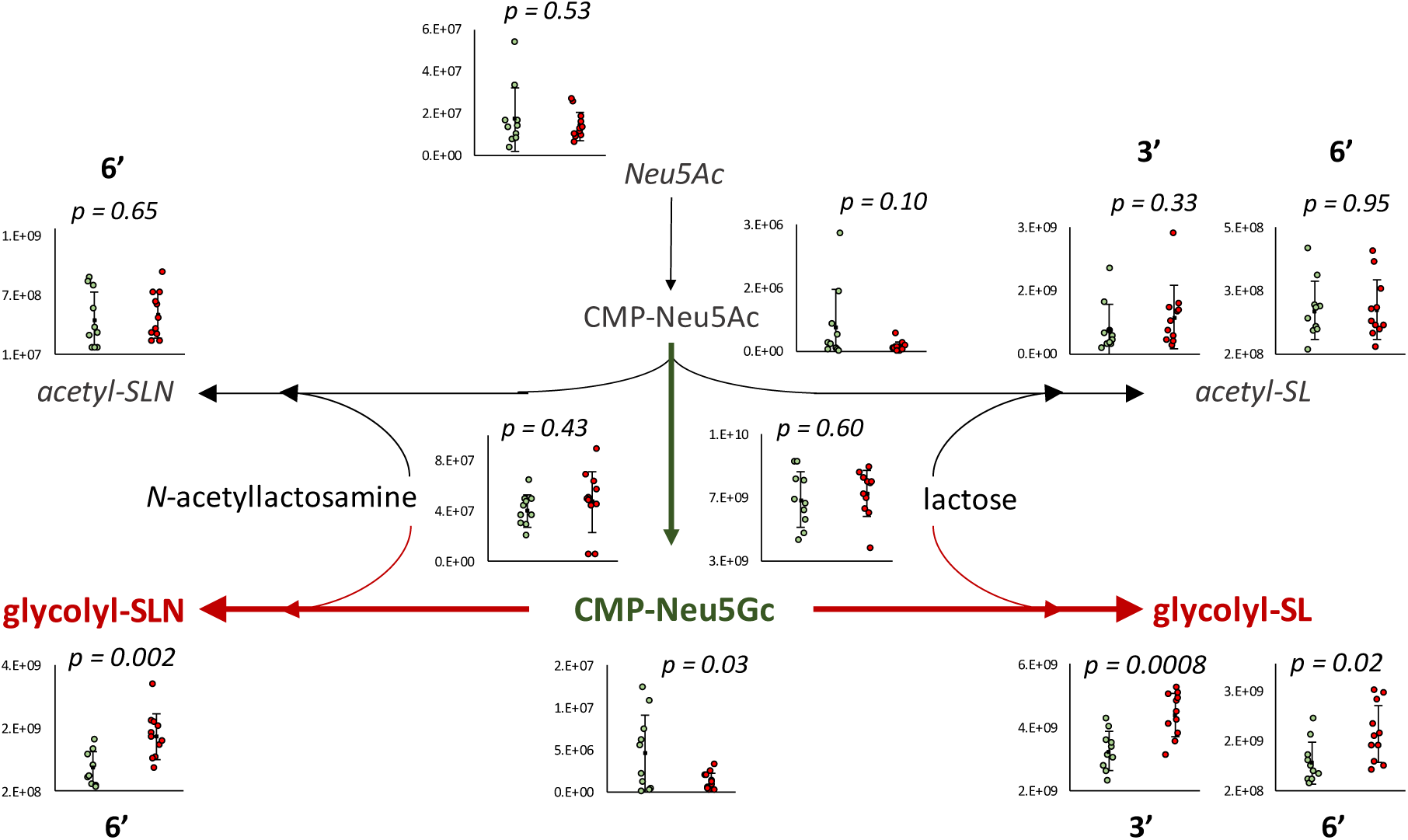
Biosynthetic pathway of sialyl-oligosaccharides with compounds observed in significantly higher concentration in the colostrum of C (green) and SC (red) ewes at T5h. 3’ and 6’ indicates the position of the sialyl group on the OS.

The peak area ratios of CMP-Neu5Gc/CMP-Neu5Ac showed that the concentration of CMP-Neu5Gc was much higher (∼20 fold on average, with large individual variations) in both C and SC groups (p = 0.017 and p = 0.002 respectively, Supplementary Figure S2), meaning that there is no clear evidence that only specific enzymatic activity of Neu5Gc sialyl transferases was altered.

#### c Bacterial population of colostrum samples

Q-PCR analysis showed that total bacterial population was stable over time and no effect of supplementation was observed, possibly due to very high inter individual variability (Table 5). Overall, in C group, total bacterial population was on average of 1.13 × 10^6^ CFU/g while a slightly higher concentration was observed in SC group (average of 1.54 × 10^6^ CFU/g colostrum).

**Table 5:** Mean and SD of total bacterial population (16S rDNA copies/g colostrum) in C or SC colostrum samples over time and p-values associated through linear mixed model.

|  |  | Time (h) |  |  |  |  | p-value |  |  |
| --- | --- | --- | --- | --- | --- | --- | --- | --- | --- |
| Group |  | 0 | 5 | 12 | 24 | 72 | Time | Supplementation | T x S |
| C | Mean | 7.38 x 10<sup>5</sup> | 8.14 x 10<sup>5</sup> | 9.68 x 10<sup>5</sup> | 2.33 x 10<sup>6</sup> | 7.86 x 10<sup>5</sup> | 0.37 | 0.45 | 0.89 |
|  | SD | 8.21 x 10<sup>5</sup> | 5.86 x 10<sup>5</sup> | 1.52 x 10<sup>6</sup> | 5.51 x 10<sup>6</sup> | 12.3 x 10<sup>5</sup> |  |  |  |
| SC | Mean | 4.35x10<sup>5</sup> | 1.01x10<sup>6</sup> | 2.57x10<sup>6</sup> | 2.42x10<sup>6</sup> | 1.29 x 10<sup>6</sup> |  |  |  |
|  | SD | 2.49 x 10<sup>5</sup> | 1.52 x 10<sup>6</sup> | 5.12 x 10<sup>6</sup> | 3.56 x 10<sup>6</sup> | 2.43 x 10<sup>6</sup> |  |  |  |

Statistical analysis on alpha diversity indices indicated no effect of supplementation whatever the indexes considered (p > 0.05), possibly because of the low number of samples considered. Indeed, higher numerical values of richness (i.e. Observed OTUs and Chao1) and evenness (Shannon index) were observed in C group (Figure 9). One outlier sample from C group (C_6) presented lower richness and evenness than the other control samples, while one outlier sample from SC group (SC_5) presented a higher richness than the rest of SC samples. Based on the visualization of the beta diversity using Bray-curtis distance and PCoA projection, no clusterization of bacterial community structure at OTU level was identified according to experimental groups (Figure S3). The colostrum sample C_2 from C group was apart from all the others, both along the first (2.82 %) and the second axes (15.9 %).

**Figure 9:**
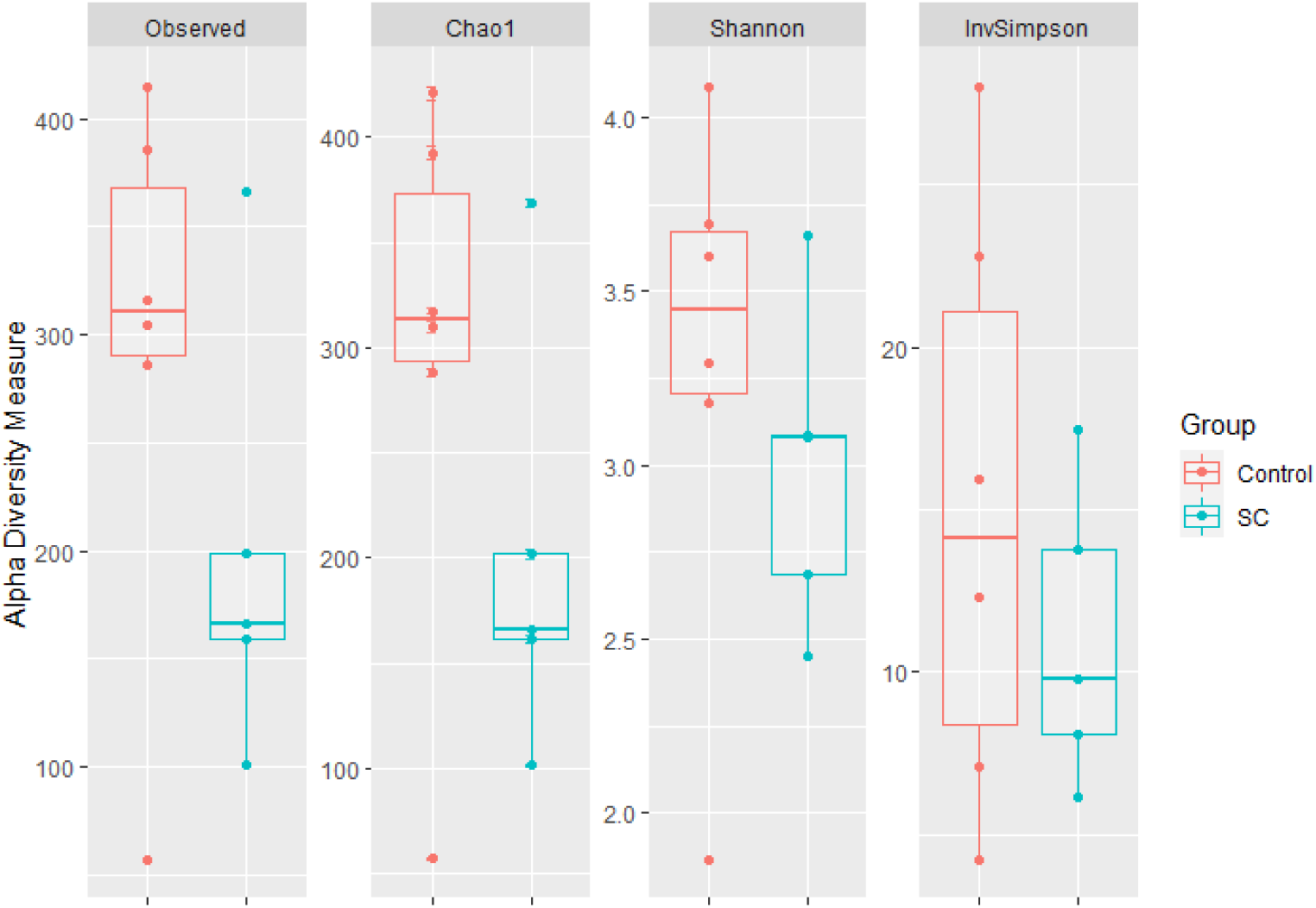
Alpha diversity indices of T0-colostrum samples from C (n = 6) and SC (n = 5) groups.

Bacterial taxonomic composition of T0-colostrum samples at the Phylum and Family levels is presented in Figure 10. At the Phylum level (Fig 10a), colostrum samples were dominated by 5 major phyla with Proteobacteria and Firmicutes being the most abundant in C (36.30 ± 26.4 % and 33.57 ± 15.98 %, respectively) and in SC samples (35.93 ± 9.61 % and 28.81 ± 9.34 %, respectively). Actinobacteria was the third phylum in terms of relative abundance in both groups (18.36 ± 10.81 % and 25.29 ± 8.94 % in C and SC group, respectively). Numerical differences were observed between groups (C vs SC) with a higher level of Proteobacteria and a lower level of Actinobacteria in C samples, but these differences were not statistically significant (p > 0.05). The previously identified unique C_2 sample was characterized by the highest abundance of Proteobacteria (72.55 %).

**Figure 10:**
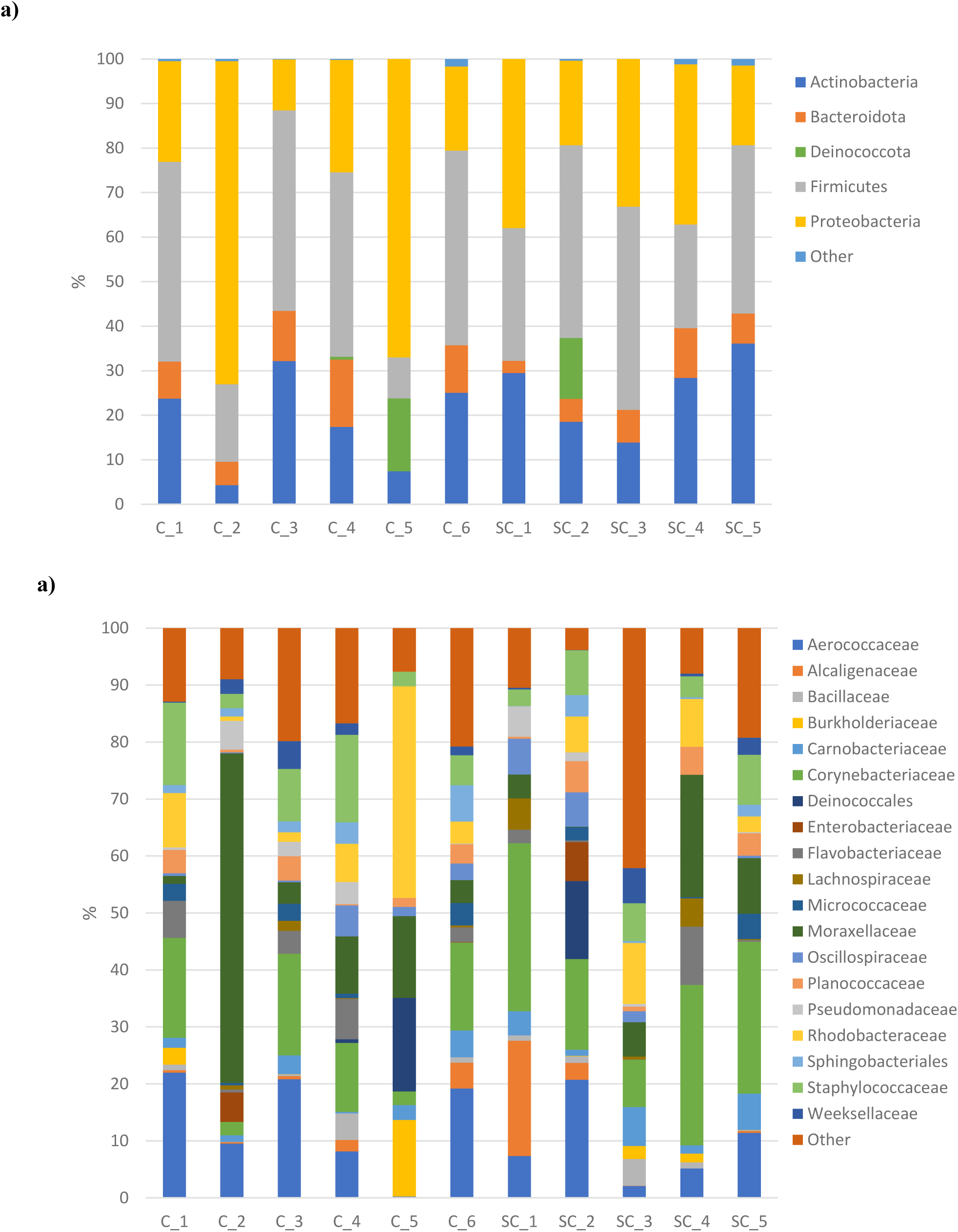
Bacterial composition of T0-colostrum samples from C or SC groups at the Phylum (a) and Family (b) levels (only Families > 1% relative abundance were represented).

At the Family level (10b), part of the bacterial diversity was linked to low abundant families (< 1 %, representing 14.47 ± 5.54 % vs 16.72 ± 15.27 % of the total family reads in C and SC groups, respectively). Aerococcaceae (13.31 ± 8.29 % vs 9.34 ± 7.21 %), Corynebacteriaceae (11.28 ± 7.19 % vs 21.7 ± 9.21 %), and Moraxellaceae (15.23 ± 21.4 % vs 8.3 ± 8.15 %) were the dominant families in C and SC groups respectively. Interestingly, a higher level of Staphylococcaceae was observed in C samples (8.22 ± 5.73 %) compared to SC group (5.96 ± 2.58 %). However, none of these observed differences reached the significance level.

## Discussion

### Colostrum composition dynamics across the first 72h post-partum

Good colostrum management at birth is recognized as an important parameter to ensure the sanitary status of the neonates, further herd performances and limit economic losses ^2^. Variations in colostrum nutrients, vitamins and minerals have been observed in several ruminant species ^3,42,43^. In the current study, the nutritional composition of colostrum changed over time and our results are in line with the literature. In ovines, variations from 0h-colostrum to milk are characterized by a slight decrease in fat content (ratio 0h-colostrum/milk ∼1-1.5), an increase in lactose (ratio ∼0.8) and a very drastic decrease in protein content (ratio ∼2-4) ^44^.

Among the most abundant and studied proteins in colostrum are immunoglobulins (Igs). The ruminant’s placental structure doesn’t allow any transfer of Igs from dam vascular system to the fetus, thus depriving newborn ruminant of antibodies at birth ^45^. Therefore, timely ingestion and absorption of colostral Igs is critical for the survival of ruminant neonates. Our results showed a drastic decrease of colostral IgG over time, an observation also seen by others ^44,46,47^. The level of IgG in colostrum varies between breeds and ranges from 17.9 to 89.3 mg/ml depending on the ovine breed ^48,49^, in line with the highest level of 79.5 mg/ml observed in the Romane ewes monitored in our study. The newborn immune system takes weeks to months to mature and become protective, and thus immune passive transfer through ingestion of IgG from colostrum is of paramount importance. Immunoglobulins improve animal growth, defenses against enteric infection by immunomodulation, mucin protein and/or modification of commensal microbial composition ^50^. Although long term health parameters were not monitored in this study, the level of Ig in the serum of lambs at birth has been associated with a better survival during the neonatal period ^6^ and delaying colostrum feeding within 12h of life has been shown to decrease the passive transfer of IgG, possibly leaving the calf more vulnerable to infections during the preweaning period ^51^. The authors also observed a delay in bacterial gut colonization, increasing the risk for pathogen colonization of the neonate. In our study, the IgG level in lamb’s serum drastically decreased with time, with a significant impact of the rearing mode. More precisely, lambs from the Cont-Art group presented 30 % less IgG in their serum after 2d than C lambs kept with their mothers, indicating a failure in the immune passive transfer. Lactoferrin is part of the high abundant protein pool mainly produced by the mammary epithelial cells, and possesses antibacterial, antifungal, antiviral and antiparasitic activities ^10,52^. It is of particular significance against *Staphylococcus aureus*, the most common mastitis-related pathogen in sheep ^53^. In young ruminants, lactoferrin significantly reduced mortality and culling rate when administered to preweaned calves ^54^ and decreased the number of days of disease, with less severe diarrhea cases in calves ^55^. Lactoferrin concentration in ruminant colostrum greatly varies among species and breeds ^46,56,57^. In ewes, Navarro et al. ^47^ measured the highest lactoferrin concentration of 0.72 mg/ml 24h after lambing, and afterwards, it decreased with time. Due to the sampling time, these values are lower than those observed in our study (average of 3.59 ± 0.2 mg/ml over the first 24h after lambing).

The metabolomics analysis performed in this study allowed us to describe, for the first time, ewe colostrum metabolites over the first 72h post-partum, with a focus on oligosaccharides (OS) composition. Our results indicated a decrease of OS concentration over time, in agreement with the literature ^58^. Sialic acid – part of OS – include N-acylneuraminic acids and their derivatives, with N-acetylneuraminic acid (Neu5Ac) and N-glycolylneuraminic acid (Neu5Gc) being the most abundant. In ruminants, great variations of OS concentrations are observed ^59,60^. In ovines, in agreement with our results, Neu5Gc is a dominant component among the colostrum sialic acids ^61^, whereas in bovine colostrum, Neu5Ac compounds are predominant with great amounts of3′-sialyllactose (3′SL) ^62^. Neu5Gc is found in variable concentrations in other ruminant species’ colostrum, depending on breed ^63^. Whereas Neu5Gc represents a great proportion of total sialic acids in bovine colostrum (~32 %), much lower levels (~6 %) are found at d90 of lactation in bovine milk ^64^. Such variations have also been observed along the lactation period in goat colostrum and milk, ranging from 40.1 to 7.4 % ^65,66^.

To our knowledge, this study is the first to describe ovine colostrum bacterial composition up to the Family level. T0-colostrum samples were collected as aseptically as possible, but contaminations from the environment, animal or humans cannot be excluded. Despite inter-individual variability, total bacterial population was very stable over time at a concentration of ~10^6^ copies of 16S rDNA gene/g. These results are close to the concentrations obtained through classical bacterial enumerations reported by Lindner et al., ^67^ and through qPCR quantification by Klein-Jöbstl et al. ^20^ on bovine colostrum (around 5 log_10_ CFU/ml and a median of 4.55 × 10^5^ copies/g, respectively). Bacterial composition of bovine colostrum and milk was described in several studies, confirming the presence of a resident microbiota in these biological fluids, however exact inoculation mechanisms remain unclear. In addition to teat skin as a potential source of inoculation, the hypothesis of an enteromammary route for colostrum and milk microbiota has been suggested in several studies in humans and ruminants ^68^. In ruminant, bacterial OTUs belonging to *Ruminococcus, Bifidobacterium* and *Peptostreptococcaceae* were observed in both milk and feces in lactating cows ^69^ and the enteromammary pathway of milk microbial inoculation has been suggested to occur through bacterial transport by immune cells ^70^. In our study, we observed a dominance of Proteobacteria, Firmicutes and Actinobacteria in T0-colostrum samples. More precisely, at the Family level, high relative abundances of Aerococcaceae, Corynebacteriaceae, Moraxellaceae and Staphylococcaceae were observed. Members among Aerococcaceae and Moraxellaceae taxa are commonly associated with mastitis, while *S. aureus* is considered one of the main species responsible for clinical mastitis in dairy ruminant ^71,72^, highlighting the presence of potential pathogens in ovine colostrum. Dominance of Proteobacteria in cow colostrum samples was observed in several studies ^19,20^, while Lima et al. ^73^ observed a dominance of Firmicutes in colostrum from both primi- and multiparous cows. This heterogeneity in bacterial composition is also observed at genus level as *Lactobacillus, Staphylococcus, Bifidobacterium* and *Akkermansia* were reported ^19^ but typical members of rumen ecosystem were also observed such as *Prevotella* and *Ruminococcaceae* ^73^. In one study, *Enhydrobacter* was found to be predominant in cow colostrum ^20^ while facultative anaerobic bacteria such as *Streptococcus, Acinetobacter, Enterobacter*, and *Corynebacterium* were observed in another work on the same animal species ^74^. The presence of anaerobic and aerobic bacteria in colostrum may participate to early gut colonization ^75^. The high heterogeneity of colostrum bacterial composition found in literature might be linked to the variability observed in the other parameters (nutrients, bioactive molecules, IgG), as it is generally reported for this type of biological fluid. No correlation could be drawn between colostrum microbiota and any of the studied parameters in the present work, due to the limited number of animals. Further studies are needed to bring more insights into potential mechanisms responsible for microbial inoculation of colostrum in the mammary gland.

### The effect of live yeast supplementation in late gestation on colostrum and hypothesis on mechanisms involved in colostrum quality improvement

To date, very few publications have addressed the interest of probiotics as a promising nutritional strategy in gestating ruminants to improve offspring health and performance through optimization of colostrum management. It has been shown that peripartum cows’ supplementation with a 2-strains cocktail didn’t improve colostrum and calves serum immunoglobulin concentrations, with only slight changes on colostrum nutrients yield ^76^, highlighting the importance of several parameters such as the administrated strain, the dose or the animal species considered. In our study, SC supplementation to the gestating ewes was shown to increase bioactive molecules in colostrum, especially the Neu5Gc containing sialylated OS and the IgG concentrations. A significant increase of serum IgG concentration for lambs in artificial rearing system is likely to result in a beneficial long term effect on lamb immunity. No SC effect was seen on colostrum composition over time in our study, in accordance with Macedo et al., ^77^ who observed no effect of a culture of *S. cerevisiae* on ewes’ colostrum nutritional composition and yield. Contrarily, a cocktail of *Bacillus licheniformis* and *B. subtilis* (BioPlus 2B) supplementation increased daily milk yield, fat and protein contents and decreased lamb mortality (mainly due to diarrhea) from 13.1% to 7.8 % ^78^. Stress hormones such as catecholamines are able to stimulate bacterial pathogens growth by enabling iron scavenging from normally inaccessible transferrin or lactoferrin ^79^. An increase in lactoferrin content is thus of interest to prevent the growth of enteropathogenic bacteria in stressed ruminants. In our study, the numerical increase of lactoferrin content in colostrum due to SC supplementation could be of interest both for the protection of the mammary gland and the neonate health.

A higher level of colostral IgG was observed in the SC group, suggesting either a higher level of IgG in the serum of the dam or a promoted IgG transfer efficiency into the mammary gland of supplemented ewes. Once suckling is achieved, the neonate’s serum immunoglobulins rise rapidly ^8^. This process of passive immunoglobulin absorption in the intestine ceases at about 24h of age and is referred to as intestinal closure ^80^. In our study, no effect of supplementation was seen in serum IgG of lambs kept with their mother. It can be hypothesized that as serum IgG of these lambs was already measured in high concentrations, the increase observed in colostral IgG in supplemented ewes was not sufficient to induce a biological and observable difference in offspring. On the contrary, when lambs had access to colostrum only for 12h before being separated from their dam and fed with milk replacer (Art groups), a significantly higher IgG concentration in the serum of lambs born from supplemented dams was observed. The risk for Failure of Passive Transfer (FTP) was also reduced in this group, as 3 lambs out of the 10 animals sampled at d2 in C group were considered as having a FTP (serum IgG concentration < 10 g/l), while it was observed for only 1 animal out of 10 in the SC group. Interestingly, the IgG level of this Art-Suppl group was similar to those observed in mothered lambs (Mot-groups) indicating a similar passive transfer of immunity. With SC supplementation to ewes, there would be thus a potential to counterbalance the negative impact of early mother separation and incomplete colostrum feeding in neonate lambs. In lambs, Igs production starts gradually several weeks after birth with detection of endogenous IgG2, IgM and IgG1 in their serum after 2, 3 and 7 weeks respectively ^81^. The significant improvement in serum IgG status in artificially fed lambs born from supplemented dams actually represents a real benefit in terms of protection against diseases during the neonatal period until the young animal starts to produce its own antibodies. The indirect effects of probiotic supplementation to the mother on offspring immunity has previously been studied by Wójcik et al. ^82^. The authors supplemented a brewer’s yeast from the 4th month of ewe gestation or after lamb birth (15 or 30 g/h, respectively) and observed an increase in specific and non-specific humoral and cellular immunity in lambs (lysozyme, Igs, blood cells activity). The effects of direct probiotic supplementation on neonate immunity have been addressed in several studies with contradictory results. In neonate calves supplemented with *Lactobacillus acidophilus* and *L. plantarum* or *L. plantarum* only, a significantly slower decrease in serum IgG was observed compared to the control group ^83^. The yeast probiotic *Saccharomyces boulardii* at 10^6^ and 10^7^ CFU/g in the supplemented feed enhanced blood IgG in lambs following a vaccine against BoHV-5 virus and *Escherichia coli*, thus enhancing the humoral immune response ^84,85^. On the contrary, supplementation of *L. plantarum* to dairy goats did not impact blood IgG, IgM, and IgA concentrations ^86^.

OS are of great interest for neonate health as they can reach the intestine and promote beneficial bacterial growth. Indeed, high levels of OS were observed in the distal jejunum and colon respectively 6 h and 12 h after colostrum ingestion by calves (~400 μg/g for 3’SL) ^15^, which was linked to an increase in mucosa-associated *Bifidobacterium* in the distal jejunum and colon ^12,51^. Overall, OS have been shown to inhibit adhesion of *Pseudomonas aeruginosa, Escherichia coli* O157:H7 and *Staphylococcus aureus* to epithelial cells ^87^ as well as ETEC strains binding to the intestinal epithelium ^16^. More precisely, Neu5Gc is a receptor for several pathogens ^88–90^ and may appear as a first line of defense to limit pathogen internalization process. Thus, the increase of Neu5Gc compounds in the colostrum of supplemented ewes may increase newborn protection against various pathogens in the neonatal period, as well as promote growth of beneficial bacterial in the intestine. In our study, a significant increase in Neu5Gc concentrations was observed up to 5 h after birth in the colostrum of supplemented ewes, with 6’-GSL and 6’-GSLN being highly enriched. The reconstruction of the biosynthesis pathway of Neu5Gc suggested a higher activity of sialyltransferase transforming CMP-Neu5Gc into Neu5Gc compounds, although it might not be the only alteration in the pathway. In a human study, a cocktail of probiotic bacteria modulated human milk oligosaccharide profile, increasing concentrations of 3’-sialyllactose and 3-fucosyllactose and decreasing concentration of 6’sialyllactose ^91^, but no study considering probiotic administration and colostrum OS composition in ruminants has been published yet.

Colostrum and milk microbiota have been shown to be affected by probiotic administration in humans. Indeed, administration of a multi-strain probiotic during the perinatal period resulted in increased lactobacilli and bifidobacteria in colostrum and milk of mothers with vaginal delivery compared to placebo ^92^. Noteworthy, mothers with caesarian sections in the probiotic group didn’t present similar bacterial increases. In addition, the same viable strain of obligate anaerobe *Bifidobacterium breve* was identified at the same time in human breast milk and both maternal and neonatal feces in one mother-child pair, demonstrating the existence of vertical mother-neonate transfer of maternal gut bacteria through breast feeding ^93^. In our study, we couldn’t identify significant differences in colostrum microbiota of the two groups, maybe due to high inter-individual variability and low number of samples. However, a higher numerical relative abundance of Corynebacteriaceae was observed in colostrum from the SC group. This taxon has been suggested to represent protection against mastitis pathogens through competition for niche adaptation by Porcellato et al., ^72^.

Currently, the exact mechanisms by which oral probiotics would affect colostrum composition remains unclear. The reasons for the observed increase of lactoferrin and colostral IgG concentrations due to SC supplementation are not yet understood; nor are those explaining the modifications of OS pathways leading to higher levels of Neu5Gc compounds in ewes’ colostrum. In a human study, Mastromarino et al. suggested that orally distributed probiotic exerted a systemic effect through the improvement of various extra intestinal conditions, such as enhancement of immune activity and modulation of systemic inflammation and metabolic disturbances ^92^. It can be hypothesized that probiotics, known to have beneficial effects on the GIT microbiota, would influence the bacterial transfer into the mammary gland and ultimately the microbial composition of colostrum and milk. According to this hypothesis, the ruminal microbial modifications expected with SC supplementation might impact lower gut microbiota, as suggested by studies of Bach et al., ^26,27^ and thus might modulate this possible microbial transfer into the mammary gland. Live yeast supplementation leads to positive impacts on rumen microbiota and health but can also exert beneficial effects beyond the rumen on oxidative status and performance ^26,31,94,95^, strengthening the systemic effect hypothesis. Further investigations are required to understand how SC supplementation could induce modifications in colostrum bioactive molecules and bacterial populations.

## Data Availability Statement

Sequencing data are available in the BioProject SRA database https://submit.ncbi.nlm.nih.gov as PRJNA732567.

## Author contribution

Conceptualization, EF, FCD; methodology, LD, JR, EF, FCD; software, LD, JR, CA; formal analysis, LD, JR, FCD; investigation, LD, JR, MS, EF, FCD; data curation, LD; writing— original draft preparation, LD, FCD; writing—review and editing, JR, MS, MC, CA, EF; supervision, LD, FCD; project administration, EF, FCD; funding acquisition, EF, FCD. All authors have read and agreed to the published version of the manuscript.

## Acknowledgments

the authors would like to thank Catherine Dhainaut and Raphaele Gresse from UMR MEDIS and Yu Jeaju from University of Guelph for their skilled technical assistance; Mickaël Bernard, Aline Cercy, Loic Gaillard, Bruno Viallard and Pierre Chirent from UE Herbipôle for their very valuable help in the animal experiment.

## Financial support

This research was partially funded by Lallemand SAS.

## Conflict of interest

LD, AC and FCD are employees of Lallemand SAS.

## Notes

### Competing Interest Statement

LD, CA and FCD are employees of Lallemand SAS

