## Supplemental tables and figures for "A live yeast supplementation to gestating ewes improves bioactive molecules composition in colostrum with no impact on its bacterial composition and beneficially affects immune status of the offspring"

**Supplementary data**

**Figure S1:** Identification of the differentially expressed compounds achieved by MS/MS. Product ion spectra of commercial standards of 6'acetyl-sialyllatose (6'-ASL) and 6'acetyl-sialyllatosamine (6'-ASLN) show product ion of Neu5Ac at m/z  290.088. The putative, glycolyl containing variants of these compounds within samples had similar MS/MS spectra, with a distinct product ion at m/z 306.0826, indicating that an additional oxygen is present on the sialyl residue, in line with Neu5Gc.

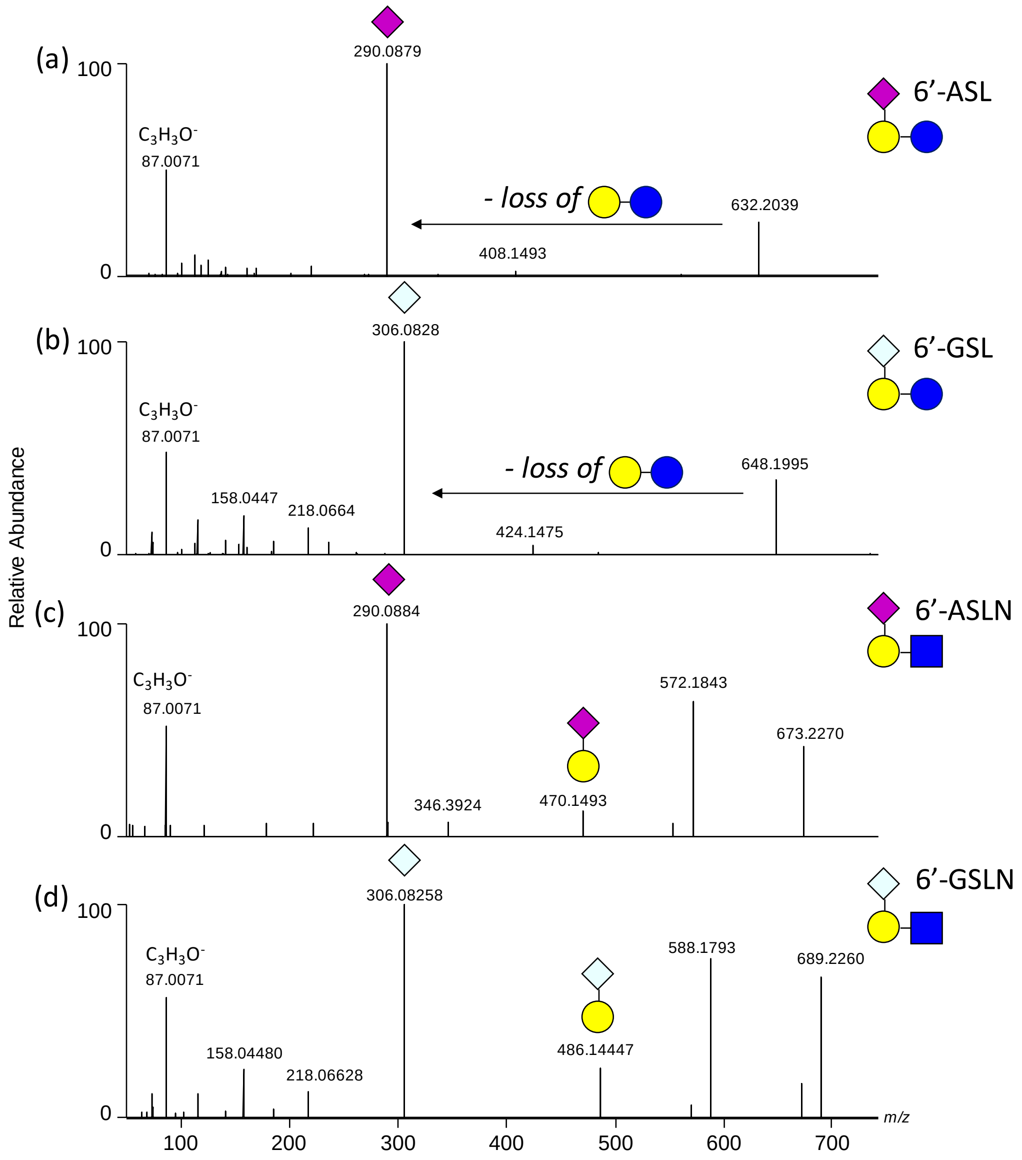

**Figure S2:** Pick area ratio of CMP-Neu5Gc/CMP-Neu5Ac in T5h colostrum from C (green) or SC (red) ewes.

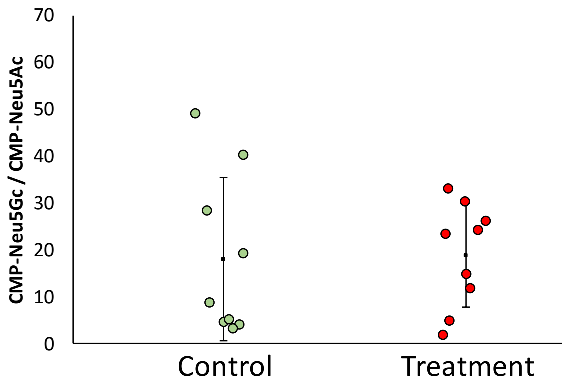


**Figure S3:** OTU PCoA plot according to Bray Curtis distance of T0-colostrum samples from C (red, n = 6) and SC (blue, n = 5) groups.

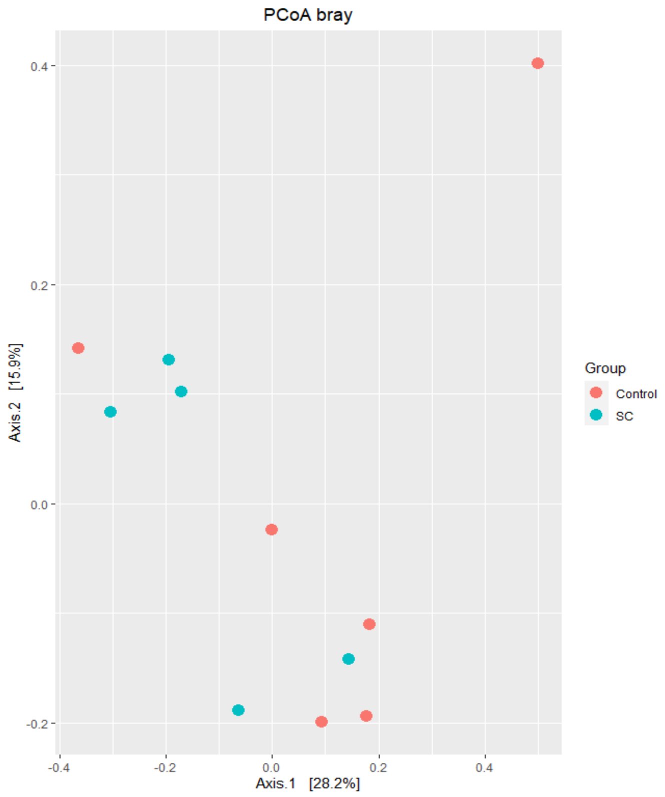

**Table S1**: Composition of the commercial concentrate.

| Ingredient | g/kg of concentrate |
|---|---|
| Corn gluten feed | 179.7 |
| Rapeseed cake | 150.0 |
| Linseed extruded supplement | 146.3 |
| Wheat bran | 127.0 |
| Barley grain | 100.0 |
| Cereal by products | 62.0 |
| Corn meal | 50.0 |
| Sugarcane molasse | 50.0 |
| Beet pulp | 50.0 |
| Wheat grain | 45.0 |
| Supplements (lime carbonate, microminerals) | 40.0 |

**Table S2:** Targeted analytes quantified by LC-HRMS.

| Abbreviation | Analyte | Formula | $[M-H]^-$ | RT (min) |
|---|---|---|---|---|
| 3'-ASL | 3'-N-acetylneuraminyl-lactose | $C_{23}H_{39}NO_{19}$ | 632.2021 | 4.42 |
| 3'-GSL | 3'-N-glycolneuraminyl-lactose | $C_{23}H_{39}NO_{20}$ | 648.1964 | 4.64 |
| 6'-ASL | 6'-N-acetylneuraminyl-lactose | $C_{23}H_{39}NO_{19}$ | 632.2021 | 4.62 |
| 6'-GSL | 6'-N-glycolneuraminyl-lactose | $C_{23}H_{39}NO_{20}$ | 648.1964 | 4.80 |
| 3'-ASLN | 3'-N-acetylneuraminyl-N-acetyllactosamine | $C_{25}H_{42}N_2O_{19}$ | 673.2285 | 4.25 |
| 3'-GSLN | 3'-N-glycolneuraminyl-N-acetyllactosamine | $C_{25}H_{42}N_2O_{20}$ | 689.2233 | 4.45 |
| 6'-ASLN | 6'-N-Acetylneuraminyl-N-acetyllactosamine | $C_{25}H_{42}N_2O_{19}$ | 673.2285 | 4.37 |
| 6'-GSLN | 6'-N-glycolneuraminyl-N-acetyllactosamine | $C_{25}H_{42}N_2O_{20}$ | 689.2233 | 4.56 |

**Table S3:** Characteristics (m/z, retention time (RT) and putative ID) of differentially expressed features from positive and negative ionization modes, at 0, 5 and 72 h after lambing at p-value < 0.01, according to t-test between SC and C groups.

| | | | | p-values | | |
|---|---|---|---|---|---|---|
| **ionization mode** | **m/z** | **RT (min)** | **putative ID** | **0h** | **5h** | **72 h** |
| positive | 157,17009 | 4,923592 | $C_9H_20N_2$ | | 0,000509 | |
| positive | 167,01244 | 11,3731 | | | 0,005154 | |
| positive | 189,13474 | 4,897401 | homo-Arg | 0,002974 | | |
| positive | 204,12311 | 3,197402 | acetylcarnitine | | 0,000311 | |
| positive | 258,11045 | 5,674051 | Glycerophosphocholine | | 0,003336 | |
| positive | 263,16775 | 1,897495 | | 0,008535 | | |

| | | | | | | |
|---|---|---|---|---|---|---|
| positive | 284,33096 | 11,29636 | noise | | 0,002655 | |
| positive | 290,08908 | 0,720561 | | | 0,000254 | |
| positive | 292,20126 | 0,642597 | | 0,007958 | 0,002638 | |
| positive | 308,09742 | 4,611503 | | | 0,003651 | |
| positive | 311,0774 | 3,67844 | | | | 0,002172 |
| positive | 364,12214 | 3,714425 | | | 0,007138 | |
| positive | 376,25871 | 0,616248 | | | 0,009907 | |
| positive | 381,07872 | 3,629511 | | | | 0,001469 |
| positive | 381,07906 | 3,976418 | | | 0,00542 | |
| positive | 382,08188 | 3,396364 | | | | 0,001992 |
| positive | 393,28529 | 0,616248 | | | 0,007706 | |
| positive | 395,29085 | 0,61621 | | 0,008622 | | |
| positive | 421,31634 | 0,616685 | | | 0,001275 | |
| positive | 422,31968 | 0,61716 | | | 0,008526 | |
| positive | 432,23713 | 0,627395 | | | 0,000612 | |
| positive | 449,34751 | 0,615718 | | 0,002977 | 0,009812 | |
| positive | 525,13056 | 4,087246 | | 0,005269 | | |
| positive | 691,23962 | 4,560937 | SLN_O | 0,004065 | 0,003 | |
| positive | 692,24285 | 4,556318 | SLN_O | 0,004142 | 0,003019 | |
| positive | 713,22131 | 4,561031 | SLN_O | 0,002521 | 0,001055 | |
| positive | 714,22439 | 4,560253 | SLN_O | 0,002953 | 0,00087 | |
| positive | 715,22636 | 4,556318 | SLN_O | 0,00335 | 0,001921 | |
| positive | 814,24268 | 4,366603 | | 0,006686 | | |
| negative | 178,81327 | 1,392473 | | 0,002988 | | |
| negative | 297,73256 | 2,043758 | | 0,000823 | | |
| negative | 316,73821 | 0,891627 | Copper diIodine | | 0,001001 | |
| negative | 318,73637 | 0,891945 | Copper diIodine | | 0,000815 | |
| negative | 322,04392 | 4,270469 | | | | 0,009128 |
| negative | 332,0833 | 2,947639 | | | 0,007637 | |
| negative | 332,08335 | 2,410796 | | | | 0,004293 |
| negative | 332,08338 | 0,399051 | | 0,000245 | | |
| negative | 332,08365 | 1,344488 | | 0,004243 | | |
| negative | 629,05825 | 4,652889 | | | | 0,008088 |
| negative | 648,19769 | 4,623225 | 3’SL+O | | 0,000774 | |
| negative | 666,14627 | 3,509148 | | | | 0,004436 |
| negative | 689,22418 | 4,564851 | 6’SLN+O | 0,00242 | 0,002257 | |
| negative | 690,22785 | 4,557011 | 13C 6SLNO | 0,001758 | 0,003294 | |
| negative | 955,29114 | 5,12 | DSL$_2$O | | 0,007588 | |
